# Typhoid toxin of *Salmonella enterica* induces ISG15 response mediating host cell survival and bacterial dissemination

**DOI:** 10.1101/2025.04.29.649991

**Authors:** Daniel S Stark, Michelle King, Angela EM Ibler, Nadia Baseer, Ellen G Vernon, Yifeng Zhang, Christopher Staples, Lilliana Radoshevich, Daniel Humphreys

## Abstract

The typhoid toxin is a secreted virulence factor of typhoidal serovars of the bacterial pathogen *Salmonella enterica* implicated in typhoid fever and chronic infections. The toxin causes a DNA damage response in human cells, characterised by cell-cycle arrest and cellular distension, resulting in cellular senescence and increased bacterial burden. To better understand host responses to typhoid toxin, we performed a transcriptomic analysis of intoxicated host cells and found that the toxin induced expression of genes relating to the type-I interferon response, including the ubiquitin-like protein ISG15. ISG15 was upregulated in a STING-dependent manner, reduced bacterial burden, and was found to be critical to host cell survival in response to the typhoid toxin and purified interferon. This highlights ISG15 as an important component of the host cell defence to the typhoid toxin.

## Introduction

The typhoid toxin is secreted by typhoidal serovars of *Salmonella enterica* such as *S.* Typhi, which causes typhoid fever, a disease responsible for 21 million cases worldwide and 200,000 deaths a year. The toxin has been implicated in causing typhoid fever symptoms (Song, Gao and Galan, 2013), but there is also evidence to suggest its activity contributes to chronic and systemic *Salmonella* infection *in vivo* (Del Bel Belluz *et al*., 2016; Miller *et al*., 2018). Chronic carriage of *Salmonella* is a major cause of the endemicity of typhoid fever and has been established by the WHO as a research priority (World Health Organization, 2018). More recently, the toxin was shown to induce an anti-inflammatory response in healthy mice, suggesting a role in immunomodulation and possibly immune evasion (Martin *et al*., 2021). The typhoid toxin is also encoded by typhoidal serovar *S.* Paratyphi A and a 1-2% subset of ∼2600 non-typhoidal *Salmonella* serovars including *S.* Javiana, which cause disease in humans and food-chain animals worldwide (Den Bakker *et al*., 2011).

The toxin is comprised of a PltB pentamer linked to PltA that is bound to CdtB, which is the toxigenic DNase 1-like subunit also found in related cytolethal distending toxins (Song, Gao and Galan, 2013). Toxigenic *Salmonella* invades gut epithelial cells and is enclosed within a *Salmonella*-containing vacuole (SCV), where it secretes the toxin. The toxin is exocytosed from the infected cell and binds to sialylated glycans on surface receptors of target host cells via PltB enabling intoxication in an autocrine or paracrine manner (Spanò, Ugalde and Galán, 2008; Song, Gao and Galan, 2013). The toxin is endocytosed and CdtB is trafficked to the host nucleus by retrograde transport, where it induces the host DNA damage response (DDR) and induces cell-cycle arrest (Guidi *et al*., 2013).

The host DDR is regulated by the apical kinases ATM (ataxia-telangiectasia mutated) and ATR (ATM and rad3-related), which phosphorylate diverse substrates to orchestrate repair and cell fate decisions including survival, apoptosis and senescence (Polo and Jackson, 2011). Canonically, ATM is recruited by double strand DNA breaks (DSBs) while ATR responds to single strand breaks (SSBs) occurring during replication stress, although they have interconnected roles. The toxin induces replication stress in S/G2 phase characterised by γH2AX localisation at the nuclear periphery and hyperphosphorylation of replication protein A (RPA), leading to ATR activation and downstream phosphorylation of P53 and CHK1 (Ibler *et al*., 2019). The toxin also induces DSBs labelled by 53BP1 in G0/G1 phase, showing that the toxin induces different host DDRs which possibly lead to diverse downstream responses.

Toxin-induced replication stress leads to a senescence-like phenotype characterised by cell distension and expression of the p53 effector p21 (Ibler *et al*., 2019; ElGhazaly *et al*., 2023). The senescent cells exhibit permanent G2 arrest and a secretory phenotype that induces secondary senescence in bystander cells in a Wnt5A, INHBA and GDF15 dependent manner (Ibler *et al*., 2019; ElGhazaly *et al*., 2023). Cells treated with conditioned media from intoxicated cells underwent senescence and became more susceptible to infection, suggesting that the toxin was creating replicative niches for *Salmonella* and facilitating infection spread. It is not known whether this senescence-like phenotype also occurs as a result of DSBs in G0/G1-arrested cells, and it is possible that there are divergent host responses to the toxin.

Here we analyse the transcriptome of intoxicated cells with the aim of uncoupling host responses to the toxin, which implicates the ubiquitin-like protein ISG15 in the host responses to pathogen-induced DNA damage. ISG15 is an interferon-stimulated protein with antiviral activity but its role during bacterial infection remains unclear and has yet to be identified as interacting with bacterial toxins. We found that ISG15 enforced host survival in the presence of interferon that was necessary to contain and counteract intracellular *Salmonella*.

## Results

### The Typhoid Toxin triggers an IFN-like response

Recombinant epitope-tagged PltB^HIS^, PltA^Myc^, and CdtB^FLAG^ typhoid toxin was purified using NiNTA chromatography as previously described (Ibler *et al*., 2019). Wild-type typhoid toxin (WT-toxin) and toxin containing CdtB-H160Q with attenuated catalytic activity (HQ-toxin) were purified simultaneously. To confirm the functionality of the purified toxins, 20 ng/ml of WT- and HQ-toxin was added to HT1080 fibroblast cells for 2 h to allow for toxin endocytosis, before removal and replacement with fresh media. Cells were fixed after 24 h and prepared for immunofluorescence analysis of the DNA damage marker *γ*H2AX (**Fig. 1A**). Using the upper-quartile of untreated cell *γ*H2AX fluorescence as a threshold, 87% of nuclei in WT-toxin treated cells were found to be positive for *γ*H2AX, compared to only 33% of HQ-toxin treated cells (**Fig. 1B**). In contrast, γH2AX levels induced by HQ-toxin were equivalent to untreated cells (**Fig. 1A, 1B**). WT-toxin treatment induced a greater *γ*H2AX response than that of 24 h continuous treatment with DNA-polymerase inhibitor aphidicolin (68% positivity), which acted as a positive control. Immunofluorescence also revealed that cell nuclei became distended upon WT-toxin treatment (**Fig. 1A**), which was consistent with observations in other studies (Haghjoo and Galán, 2004; Spanò, Ugalde and Galán, 2008; Guidi *et al*., 2013; Ibler *et al*., 2019).

**Fig. 1.**
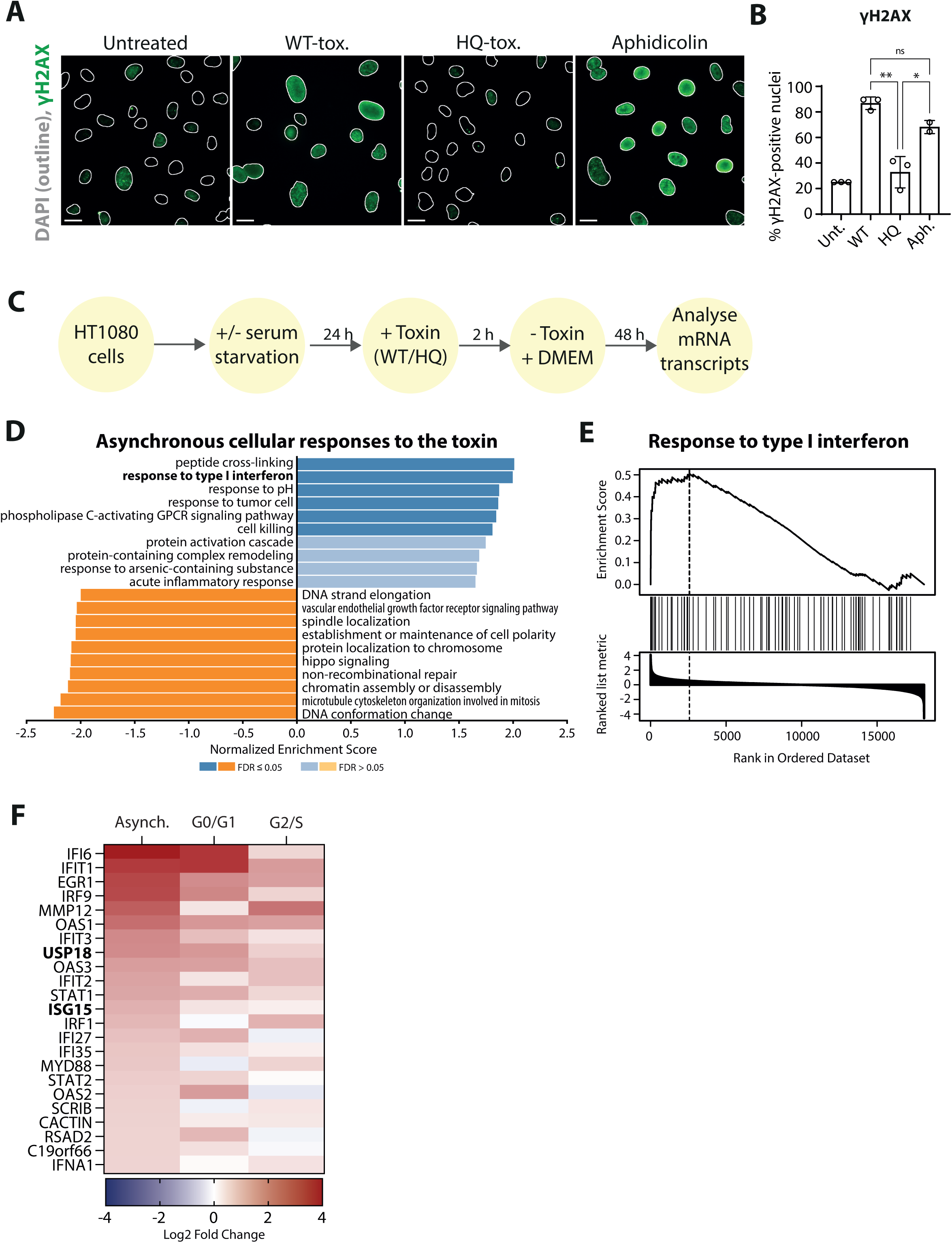
The typhoid toxin induces a type-I IFN-like response. **A**) HT1080 cells treated with 20ng/ml WT- and HQ-toxin assayed by immunofluorescence for γH2AX (green), with 20 μM aphidicolin used as a positive control. DAPI-stained nuclei indicated by white outlines. Scale bars are 20 μm. **B**) Quantification of A. Nuclei were counted as positive for γH2AX if greater than the upper quartile of untreated intensity. Each circle is from an independent experiment consisting of three replicates. Ordinary one-way ANOVA was used with Tukey’s multiple comparison test. Asterisks indicate significance *p<0.05, **p<0.01. Bars indicate mean and error bars indicate SD. **C**) Workflow for microarray analysis of differential gene expression between HT1080 cells treated for 48 h with WT- and HQ-toxin. **D**) Bar chart showing the top 10 positively and negatively enriched biological processes found using gene set enrichment analysis (GSEA) of 19460 genes analysed by microarray, compared between WT- and HQ-toxin treatment at 48 h in complete media. **E**) GSEA curve for the ‘response to type-I IFN’ biological process found in D, with the dashed line indicating the leading-edge cut-off. **F**) Heatmap showing differential gene expression of the 23 leading edge genes from E. Gene expression changes are also shown from the G0/G1 responses to the toxin, and G2/S responses to the toxin, which are shown in **Supp. Fig. 1C-E**.

Host responses to the toxin were investigated by analysing the transcriptome of intoxicated HT1080 fibroblast cells. Cells were pulsed for 2h with WT- or HQ-toxin and incubated for 48h before extraction of cellular RNA, which was analysed using a Clariom^TM^S transcriptome profiling microarray (experimental pipeline in **Fig. 1C**). 19460 analysed genes were submitted for gene set enrichment analysis using the Web-based Gene Set Analysis Toolkit (Web-Gestalt 2019) which identified positively and negatively enriched biological processes between WT- and HQ-toxin treatment (**Fig. 1D-E**). Of these processes, the response to type-I interferon was found to be positively enriched in WT-toxin treated cells compared to HQ-toxin, with 23 leading edge genes including 2 ISG transcription factors (IRF-9 and STAT1) and 10 ISGs including IFIT-1, −2 and −3, OAS-1 and −3, EGR1, RSAD2, IFI6, USP18 and ISG15 (**Fig. 1F**). This finding was consistent with studies that have shown the related bacterial genotoxin *E. coli* CDT induces a type-I interferon response in a DNA damage-dependent manner (Pons *et al*., 2021).

The enrichment of the type-I interferon response occurred following WT-toxin induced damage in asynchronous cells, meaning that toxin-dependent differential gene expression could be caused by toxin-induced RPA-labelled SSBs in G2/S phase or 53BP1-labelled DSBs in G0/G1 (**Supp. Fig. 1A, 1B**) (Ibler *et al*., 2019). Serum-starvation prevents entry into S phase, locking cells in G0/G1, which therefore inhibits toxin-induced replication stress (Ibler *et al*., 2019). In an attempt to uncouple the transcriptomic responses to these two toxin-induced types of damage, transcriptome changes were analysed following WT- and HQ-toxin treatment in the presence (10%) or absence (0%) of serum. The type-I interferon response was the most positively enriched biological process between WT- and HQ-toxin treatment in the absence of serum, with 23 leading edge genes (**Supp. Fig. 1C-D**).

Toxin-induced DSBs are permissive in the absence of serum (WT-tox., 0% serum) while SSBs in S/G2 are only permissive in the presence of serum (WT-tox., 10% serum). The major differences between the presence and absence of serum in WT-intoxicated cells were in cell cycle and cell metabolic processes, most likely due to serum-starvation blocking DNA replication and the supply of nutrients. Interestingly, the type-I IFN response as a process was not enriched, suggesting it was more likely to be a response to damage in G0/G1 phase rather than S/G2 phase (**Supp. Fig. 1E**). However, when analysing expression levels of individual ISGs, the type-I IFN response in serum-starved cells was particularly evident for ISGs such as IFI6 and IFIT1 that were observed in G0/G1 but not for ISGs such as ISG15 and RSAD2, which were up-regulated in asynchronous replication-competent cells in the presence of serum (**Fig. 1F**). This suggests divergent mechanisms underlying ISG regulation.

The differential expression of 11 of these type-I IFN genes were further examined by quantifying mRNA levels after 24h of toxin treatment using qRT-PCR. Of these, a significant increase in mRNA between WT- and HQ-toxin treatment was only seen in IFIT1, IFIT2, IFIT3, IRF-9, and ISG15 (**Fig 2A, Supp. Fig. 2**). The toxin-dependent upregulation of ISG15 in both G2/S and G0/G1 phase was particularly interesting, as ISG15 has been shown to have roles in the DDR as well as innate immune responses to bacterial invasion (Radoshevich *et al*., 2015; Raso *et al*., 2020). ISG15 is a ubiquitin-like protein and has been shown to act as a covalent adduct to other proteins throughout the cell, as well as acting alone both intra- and extracellularly (Morales and Lenschow, 2013; Perng and Lenschow, 2018). As well as ISG15 upregulation, we also observed toxin-dependent expression of USP18 (**Fig. 1F**), which catalyses the uncoupling of ISG15 from ISGylated proteins (Malakhov *et al*., 2002). Interestingly, we did not observe differential expression of any of the ISG15 ligases, such as UBE1L, which are required for ISGylation to occur (Yuan and Krug, 2001).

**Fig. 2.**
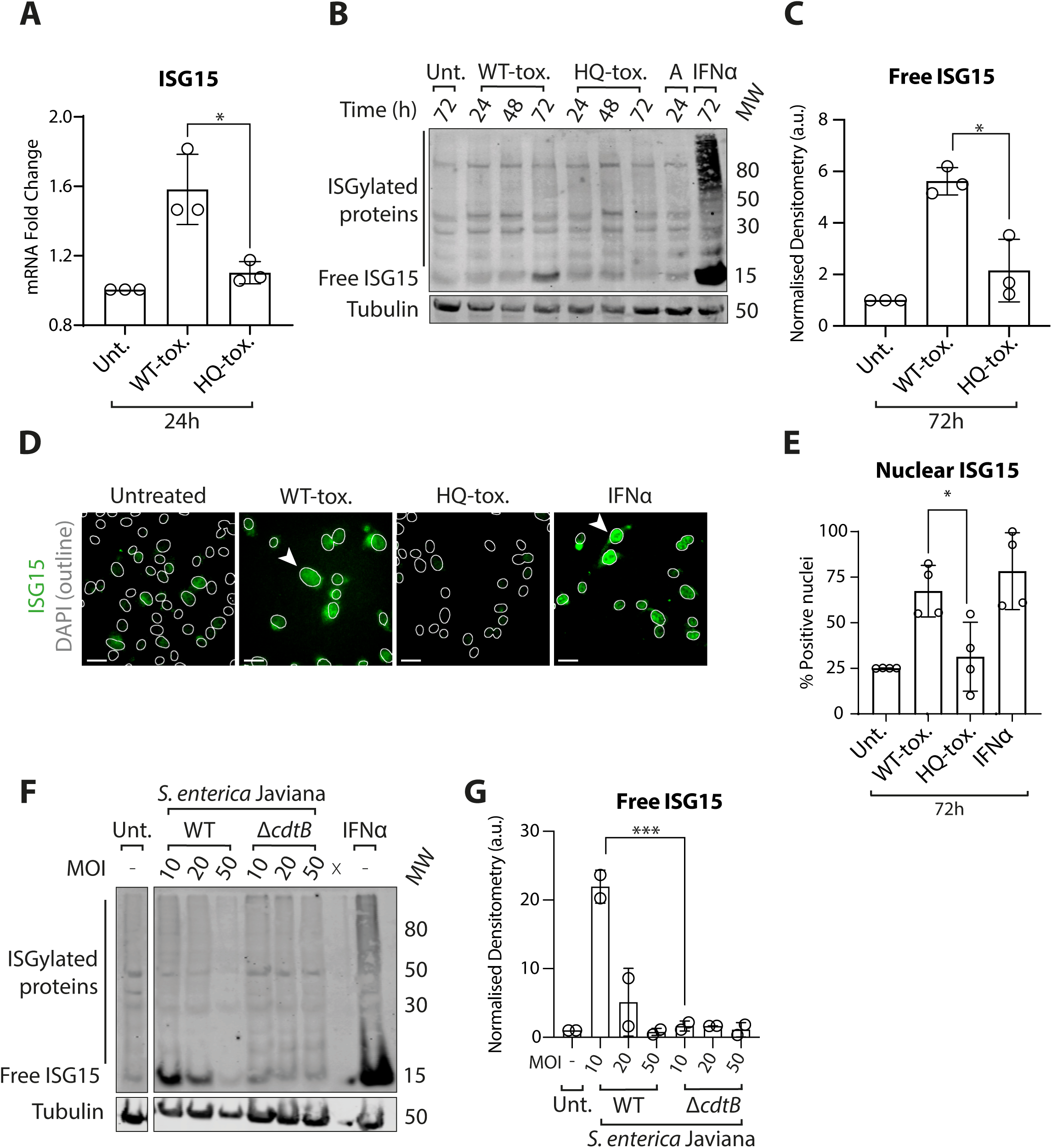
ISG15 is upregulated in response to the typhoid toxin. **A**) Quantification of ISG15 mRNA levels in untreated, WT-toxin treated, and HQ-toxin treated HT1080s cells. Each circle is an individual RNA sample with three technical replicates. WT- and HQ-toxin conditions compared using an unpaired t-test. **B**) Immunoblots of ISG15 protein levels in HT1080s 24 h, 48 h and 72 h after either treatment with WT-toxin, HQ-toxin or purified IFNα. MW indicated as kDa, right. **C**) Quantification of B showing densitometry analysis of the 15 kDa ISG15 band at 72h post-intoxication relative to tubulin and normalised to untreated (3 independent immunoblots). WT- and HQ-toxin conditions compared using an unpaired t-test. **D**) Immunofluorescence of ISG15 (green) in response to WT-toxin, HQ-toxin or purified interferon α in HT1080 cells. DAPI-stained nuclei indicated by white outlines. White arrows indicate cells with enriched nuclear ISG15. Scale bars are 20 μm. **E**) Quantification of ISG15-positive nuclei in D. Each circle is an independent experiment with three technical replicates. WT- and HQ-toxin conditions compared using an unpaired t-test. **F**) Immunoblot of ISG15 protein levels in HT1080s 144 h after 30 min infection with MOI 10, 20 or 50 of *S*. Javiana^WT^ or *S*. Javiana^ΔcdtB^. MW indicated as kDa, right. **G**) Densitometry analysis of the 15 kDa ISG15 band relative to tubulin from F and normalised to untreated (circles represent independent immunoblots). A one-way ANOVA with Šidák’s multiple comparisons test was used to measure significance relative to the untreated control, with only significant comparisons shown. Bars represent mean, error bars indicate SD, and asterisks indicate significance where *p<0.05, **p<0.01, ***p<0.001 and ****p<0.0001.

We examined ISG15 protein levels in intoxicated cells, which were significantly increased compared to both untreated and HQ-toxin treated cells, with the strongest response seen between 48 and 72h post-intoxication (**Fig. 2B-C**). As a positive control, sustained 72 h treatment with purified IFNα triggered upregulation of ISG15. IFNα-treatment also induced a smear of high molecular weight ISG15-conjugated (ISGylated) proteins which was not visible following intoxication, where only free ISG15 was observed. No toxin induced ISGylation was observed in over exposed blots (data not shown). Using immunofluorescence, we observed that ISG15 localised throughout the cell but was often enriched in the nucleus, suggesting it could be modulating the function of nuclear proteins (**Fig. 2D, 2E,** white arrows indicate enriched nuclear ISG15). ISG15 was significantly upregulated in the nucleus in WT-toxin treated cells compared to untreated and HQ-toxin treated cells.

We also examined ISG15 protein levels following infection with the toxigenic non-typhoidal *Salmonella enterica* serovar Javiana, which has previously been shown to induce a similar phenotype to that of purified toxin (Miller *et al*., 2018; Ibler *et al*., 2019; ElGhazaly *et al*., 2023). Infection induced a potent ISG15 response in a CdtB-dependent manner, but interestingly this response was not seen until 144h (6 days) post-infection (**Fig. 2F-G**). The difference in time could be attributed to the time taken for intracellular *Salmonella* to express and secrete the toxin before sufficient DDRs triggered IFN-like responses.

### The toxin causes an IFN-like response in a type-I IFN independent and STING dependent manner

ISGs are canonically regulated by IFNs binding to cognate receptors and activating a downstream JAK/STAT signalling pathway, triggering formation of heteromeric signalling complexes that bind to ISREs in ISG promoter sequences (Boxx & Cheng, 2016). Interestingly, toxin dependent ISG upregulation was only accompanied by a modest upregulation of IFNA1, with no significant differential expression of any other IFN genes. Furthermore, our attempts to validate this finding by qRT-PCR were inconclusive (**Supp. Fig. 2**). Therefore, ISG15 expression was examined following type-I IFN inhibition by B18R, which is encoded by vaccinia virus and competitively binds to IFN, thus inhibiting downstream signalling of IFN (Alcamí, Symons and Smith, 2000; Radoshevich *et al*., 2015). As expected, B18R abolished ISG15 upon addition of IFN*α* (**Fig. 3A**). However, whilst there was a reduction in ISG15, B18R did not abolish WT-toxin dependent ISG15 upregulation, suggesting that the toxin was upregulating ISG15 via both type-I IFN-dependent and - independent pathways.

**Fig. 3.**
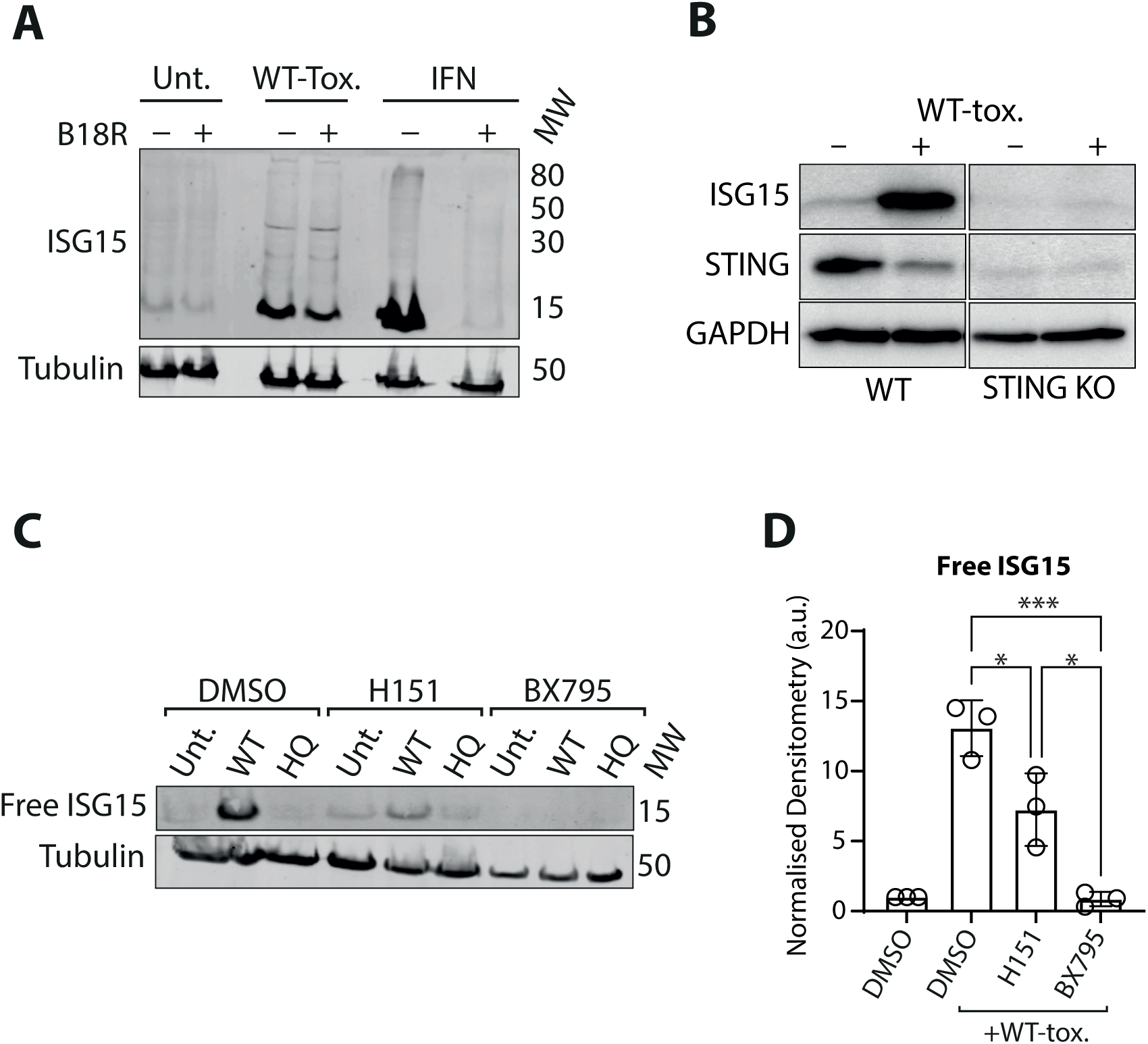
Toxin-induced ISG15 is not reliant on type-I IFN but is STING-dependent. **A**) Immunoblot of ISG15 protein levels in HT1080s 72 h after treatment with WT-toxin or purified IFNα in the presence and absence of the type-I IFN inhibitor B18R. MW indicated as kDa, right. **B**) Immunoblot of ISG15 and STING protein levels in untreated and WT-toxin treated wild-type and STING knockout HaCaT cells. **C**) Immunoblots of ISG15 protein levels in HT1080s following 72h treatment with STING inhibitor H151 or TBK1 inhibitor BX795, and treatment with WT- or HQ-toxin. MW indicated as kDa, right. **D**) Quantification of C using densitometry analysis (ImageStudio) of the 15 kDa ISG15 band intensity relative to tubulin and normalised to untreated (three independent immunoblots). One way ANOVA was used with Tukey’s multiple comparison test. Asterisks indicate significance where *p<0.05, **p<0.01, ***p<0.001 and ****p<0.0001.

We next sought to determine the mechanism by which the toxin was inducing ISG15 expression. Other studies have shown that replication stress causes cytosolic leakage of fragments of damaged DNA, activating cytosolic DNA sensors such as cGAS and STING which trigger immune responses (Wolf *et al*., 2016; Kreienkamp *et al*., 2018). We therefore explored the role of STING in response to the toxin. The toxin-dependent ISG15 response was abolished in STING-KO U2OS cells compared to wild type cells (**Fig. 3B**). ISG15 was also assayed following inhibition of STING and TBK1 by respective small molecule inhibitors H151 and BX795. Continual inhibition by either inhibitor for 72h following intoxication significantly reduced toxin induced ISG15, and indeed TBK1 inhibition by BX795 abrogated the toxin induced ISG15 response entirely (**Fig. 3C, 3D**).

### ISG15 is part of the host-defence mechanism

We hypothesised that ISG15 was upregulated as part of the host-defensive response against *Salmonella* infection and subsequent intoxication by the toxin. HT1080s were pre-treated with 72 h of continuous IFNα treatment to strongly upregulate ISG15 before infection with *S.* Javiana^WT^ or *S.* Javiana^ΔcdtB^. *Salmonella* invasion was assayed 24 h post-infection by culturing whole cell lysates on LB agar. Interestingly there was no significant difference between *S.* Javiana^WT^ CFUs from pre-treated and untreated cells (**Supp. Fig. 3A**), possibly because of the toxin-induced presence of ISG15 in both cases (**Fig. 2F-G**). However, intracellular *S.* Javiana^ΔcdtB^, which does not elicit ISG15-signalling (**Fig. 2F-G**), was significantly increased in cells without IFNα pre-treatment (**Supp. Fig. 3B**).

IFN upregulates a large array of different ISGs, meaning that the differences seen in **Supp. Fig. 3A** cannot be attributed to ISG15 alone. To uncouple the role of ISG15 from other ISGs, HT1080s were transfected with a mammalian expression vector encoding a 5HA-tagged ISG15 (pCAGGs-5HA-mISG15) which expressed HA-tagged ISG15 for up to 144 h post-transfection, (**Supp. Fig. 3B**). As before, there was no significant difference between 5HA-ISG15 and empty vector transfection in *S.* Javiana^WT^ CFU counts (**Supp. Fig. 3C**), but there was a reduction in *S.* Javiana^ΔcdtB^ CFUs following HA-ISG15 transfection (**Supp. Fig. 3C**). This further suggested that ISG15 overexpression caused a decrease in *Salmonella* CFUs indicating that ISG15 was counteracting infection. However, transfecting the cells with the vector was sufficient to induce expression of endogenous ISG15. Therefore, to analyse the role of ISG15 in the host defence response, we instead made use of a stable ISG15-/-A549 cell line (A549^ISG15-/-^).

Infection of A549^WT^ and A549^ISG15-/-^ cells with *S.* Javiana^WT^ revealed a higher bacterial burden at 24h in A549^ISG15-/-^ cells when analysed using immunofluorescence (**Fig. 4A**). This was corroborated by an increase in *S.* Javiana^WT^ CFUs in A549^ISG15-/-^ cells compared to A549^WT^ 48h post-infection (**Fig. 4B**). Addition of IFNα to induce ISG15 expression in A549^WT^ cells did not appear to impact bacterial burden but interestingly caused significant host cell loss when added to A549^ISG15-/-^ cells (**Fig. 4A**). A549^ISG15-/-^ cell loss in the presence of IFNα was coincident with a significant decrease in *S.* Javiana^WT^ CFUs (**Fig. 4B**). To ensure that the phenotype was not dependent on cell-type, we performed the same experiment in a ISG15-/-U2OS cell line (U2OS^ISG15-/-^). The increase in *S.* Javiana^WT^ burden seen in infected A549^ISG15-/-^ was more modest and not statistically significant in U2OS^ISG15-/-^ cells when compared to U2OS^WT^ cells, and *S.* Javiana^WT^ CFUs were in general found to be much lower in U2OS cells compared to A549 cells. However, in a similar manner to A549 cells, IFNα almost completely abolished infection in U2OS^ISG15-/-^ cells (**Fig. 4C**). To confirm that this was due to host cell death, we performed MTT assays in A549 (**Fig. 4D**) and U2OS (**Fig. 4E**) cells, confirming in both cell types that addition of IFNα causes significant host cell death in ISG15-/-cells compared to wild-type at 48h.

**Fig. 4.**
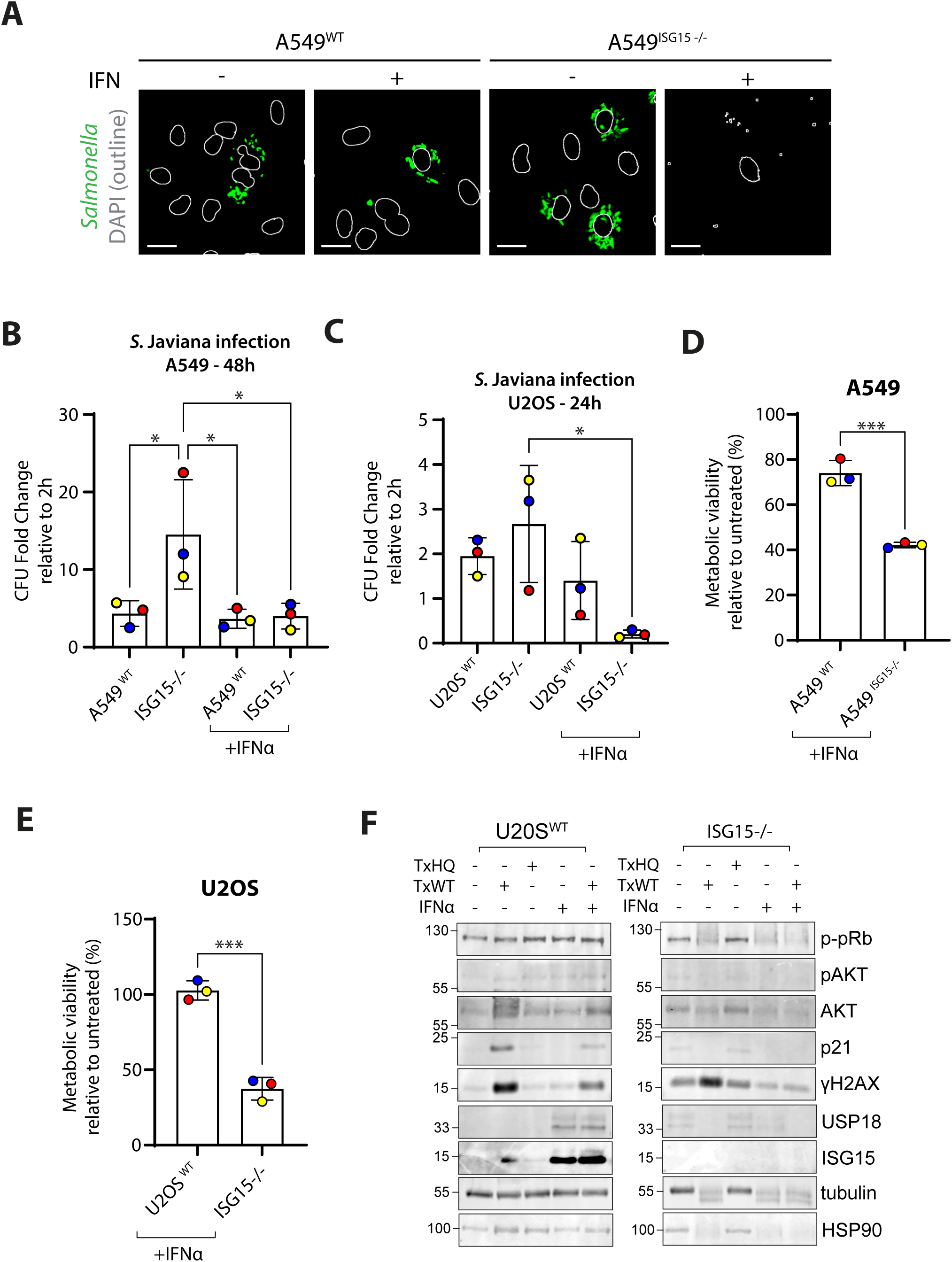
ISG15 inhibits *Salmonella* infection and is essential for cell survival in response to IFN. **A)** Immunofluorescence of *S*. Javiana^WT^ (green) in infected wild-type and ISG15-/-A549 cells with either mock or purified IFNα treatment at 48 h. DAPI-stained nuclei indicated by white outlines. Scale bars are 20 μm. **B**) *S*. Javiana^WT^ CFUs 48 h after infection in wild-type and ISG15-/-A549 cells with either mock or purified IFNα treatment. **C**) *S*. Javiana^WT^ CFUs 24 h after infection in wild-type and ISG15-/-U2OS cells with either mock or purified IFNα treatment. **D**) Cell viability determined by MTT assay in wild-type and ISG15-/-A549 cells with purified IFNα treatment. Viability is shown as a percentage of metabolic viability in the mock treatment controls. **E**) As D, but in U2OS cells. **F**) Immunoblot of ISG15, USP18, p21, γH2AX, phospho-Rb, Akt and phospho-Akt protein levels in wild-type and ISG15-/-U2OS cells after 48 h with either mock treatment, WT-toxin, HQ-toxin or purified IFNα treatment. Circles indicate separate experiments. Bars represent mean, error bars indicate SD, and asterisks indicate significance where *p<0.05, **p<0.01, ***p<0.001 and ****p<0.0001.

We hypothesised that IFNα treatment was triggering apoptosis in the absence of ISG15, which we investigated by immunoblotting control and ISG15-deficient cells. In U2OS cells. U2OS^WT^ and U2OS^ISG15-/-^ cells were treated with either typhoid toxin or IFNα, or both in combination before immunoblotting at 48h (**Fig. 4F**). Relative to untreated (-) or HQ-Tox control U2OS^WT^ cells, WT-Tox induced DDRs marked by γH2AX and elevated p21 indicative of cell-cycle arrest, which suggests entry into senescence and resistance to apoptotic signalling (Kumari and Jat, 2021). WT-Tox also induced a modest increase in ISG15, which significantly increased in the presence of IFNα, either alone or in combination with WT-Tox. USP18 expression was independent of ISG15 and required robust signalling induced by IFNα, which did not induce γH2AX or p21 signalling in isolation. Phosphorylation of the survival kinase AKT (pAKT) has been shown to promote cell survival (Brunet *et al*., 1999) and loss of pAKT has been observed in apoptotic ISG15-deficient macrophages following treatment with IFNα (Waqas *et al*., 2022). In control U2OS^WT^ cells, pAKT was observed in all conditions indicating activation of cell survival pathways. In contrast, U2OS^ISG15-/-^ cells displayed several indicators of apoptosis in the presence of either WT-Tox or IFNα: we found degradation of loading controls tubulin and HSP90, which was suggestive of caspase-mediated proteolysis. Cell death was consistent with the observation that WT-Tox failed to induce p21 senescence signalling, which is associated with pro-survival (Kumari and Jat, 2021), despite γH2AX-marked DDRs. Perhaps of most significance was the loss of pAKT signalling in the presence of WT-Tox or IFNα in U2OS^ISG15-/-^ cells. In contrast, cell death in A549 cells was not due to loss of pAKT, which was observed in the presence of IFNα in both parental A549^WT^ and A549^ISG15-/-^ cells (**Supp. Fig. 3D**). Instead a separate mechanism was apparent as ISG15-dependent expression of USP18 in A549 cells was necessary for mediating cell survival in IFNα-treated cells (**Supp. Fig. 3D-F**). Thus, ISG15 is required for pro-survival signalling in the presence of typhoid toxin or IFNα, which indicates why ISG15-deficient host cells have increased susceptibility to cell death.

## Discussion

Here, we have performed an unbiased analysis of host responses to the typhoid toxin by identifying the toxin-dependent transcriptome. By doing this, it was found that a type-I IFN like response is induced in a toxin-dependent manner. Through validation of the microarray data, ISG15 was identified as a regulator of host cell survival following *Salmonella* infection and intoxication by the typhoid toxin.

We observed that the toxin induced ISG15 expression but did not appear to be causing ISGylation of other proteins. IFN-treatment resulted in an intense smear of ISGylated proteins in cell lysates, but the same was not seen in intoxicated cells. ISGylation machinery (UBE1L, UbcH6/8 and HERC5) was not upregulated in a toxin-dependent manner in the microarray data. This suggests that toxin-induced ISG15 is free and performs a role that does not involve conjugation to other proteins. This has been seen in epithelial cells in response to *Chlamydia* infection, where ISG15 expression was found to dampen inflammatory responses and reduce bacterial burden (Wu *et al*., 2024). ISG15 expression in this scenario was also found to be independent of the Type-I IFN signalling pathway, but dependent on STING, TBK1 and IRF3, much like our observations of the ISG15 response to the typhoid toxin. Free ISG15 has also been shown to accelerate replication fork progression by non-covalent interaction with PCNA, thus inducing chromosomal breakage (Raso *et al*., 2020). It is possible that toxin-induced ISG15 exacerbates replication stress caused by the DNA damage response, leading to increased genomic instability and senescence. A positive feedback loop such as this may explain how the typhoid toxin triggers a potent DDR despite having attenuated nuclease activity compared to other DNAses (Elwell *et al*., 2001).

It was found that STING inhibition reduced toxin dependent ISG15 induction. This is consistent with findings with other CDTs, which showed activation of a type-I IFN response in a cGAS-STING dependent manner (Pons *et al*., 2021). Activation of STING canonically leads to upregulation of type-I IFNs via phosphorylation of TBK1 and IRF3. However, studies have shown that STING can be non-canonically activated by a signalling process involving ATM, P53 and IFI16, leading to NFκB signalling in DNA damage conditions (Dunphy *et al*., 2018).

ISG15 knockout increased *Salmonella* CFUs 48h post-infection, suggesting that ISG15 is performing a host defensive role. ISG15 has previously been shown to have no effect on *Salmonella* Typhimurium invasion 4 h post-infection (Radoshevich *et al*., 2015). However, these experiments were performed in mouse embryonic fibroblasts and not human cells as this study. Also, it may take a longer period (>4h) for ISG15 to exert its role. Furthermore, ISG15 knockout significantly increased *Salmonella* CFUs in A549 cells, but was not significant in U2OS cells, showing that the effects of ISG15 on *Salmonella* infection are cell type specific. The exact mechanism by which ISG15 is protecting the host against *Salmonella* infection is uncertain.

Increased ISGylation of Golgi and ER proteins promotes secretion of cytokines that counteract *Listeria* infection (Radoshevich *et al*., 2015; Zhang *et al*., 2019). However as previously discussed, there is little evidence of toxin induced ISGylation. It is possible that ISG15 may be preventing bacterial growth via modulation of P53. Studies have shown that ISG15 levels are dynamically linked to P53 activation (Park *et al*., 2016). P53 has been shown to suppress cell metabolism and thus inhibit growth of *Chlamydia*, another intracellular bacterium (Siegl *et al*., 2014). It is possible that ISG15 is increasing P53 levels and thus preventing *Salmonella* replication in a similar manner. However, further work will be needed to determine how ISG15 is exerting an antibacterial role in response to *Salmonella* infection.

ISG15 was found to be critical to host survival in response to both the typhoid toxin and purified IFN. In humans, ISG15 deficiency has been associated with IFN-I induced auto-inflammation and enhanced mycobacterial infection (Zhang *et al*., 2015). ISG15 was found to stabilise and enable the accumulation of USP18, which regulates type-I IFN signalling. Deregulation of ISG15 and USP18 has been shown to trigger cell cycle arrest via activity of the S-phase kinase associated protein 2 (SKP2) (Vuillier *et al*., 2019). This may explain why ISG15 knockout results in cell death in response to typhoid toxin or IFN.

We illuminate the host response to the typhoid toxin by an unbiased transcriptomic response that uncovers a novel host response to the typhoid toxin. We have identified ISG15 as a regulator of bacterial burden and host survival in response to *Salmonella* infection. This contributes to our understanding of the functional role of the toxin in *Salmonella* pathogenesis and presents a novel role for ISG15 in response to a bacterial genotoxin.

## Materials & Methods

### Purification of recombinant typhoid toxin

Recombinant wild-type (WT) and H160Q (HQ) typhoid toxin was purified as previously described (Ibler *et al*., 2019). Briefly, the T7 expression vector encoding pETDuet1-pltB-HIS/pltA-MYC/cdtB-FLAG was transformed into chemically competent BL21 Rosetta *E*. coli and toxin expression induced by addition of 0.1 mM Isopropyl β-D-1-thiogalactopyranoside (IPTG). Toxin was purified by addition of bacterial lysate to NiNTA agarose beads (Jena Bioscience, AC-501-25) and elution with Tris-buffered saline pH8 supplemented with 250mM imidazole. Toxin preparations were kept at −80°C with 20% glycerol.

Unless otherwise stated, 20 ng/ml WT- or HQ-toxin was added in culture media to cells for 2h. Cells were then washed 3x with sterile PBS and chased with fresh complete growth media.

### Mammalian cell culture

The A549 and U2OS wild-type and ISG15-/-cell lines were kindly donated by the laboratory of Lilliana Radoshevich at the University of Iowa. ISG15-/-cell lines were prepared using a CRISPR/Cas9 approach. The target sequences were designed using www.benchling.com and oligonucleotides synthesised via Integrated DNA Technologies (www.IDT.com). These were cloned into the following plasmids:

pX330-U6-Chimeric_BB-CBh-hSpCas9 (#42230, Addgene)

pSpCas9(BB)-2A-GFP (#48138, Addgene)

Both plasmids were gifted by Feng Zhang (Department of Biological Engineering, MIT). The cells were co-transfected with ISG15-targeting plasmids along with FU gene® HD transfection reagent (BD Science) then GFP positive cells were isolated and plated in 96-well plates for single-clone selection using Becton Dickinson FACS Aria Fusion (BD Science). Transfected cells were challenged with IFNα and subsequent lack of ISG15 expression confirmed through Western blotting. Additionally, knock out of ISG15 was further confirmed by genomic PCR and next-generation sequencing of the ISG15 amplicon.

Primers - ISG15 NCBI FWD 5’ gtgtgcctcaggcttataatagg 3’

ISG15 NCBI REV5’ cggccatttctctttacaacagcc 3’

Both primers were synthesised by Integrated DNA Technologies.

HT1080 human fibrosarcoma cells were maintained in high glucose Dulbecco’s Modified Eagle’s Medium (DMEM; Sigma Aldrich, D6546). A549 and U2OS cells were maintained in DMEM Glutamax (Gibco # 31966-021). For all cells, media was supplemented with 10% foetal bovine serum (FBS) (Sigma Aldrich, F7524), 10 U/ml Penicillin/Streptomycin (Gibco, 11548876), 50 μg/ml Kanamycin sulphate (BioBasic, KB0286) and 2 mM L-glutamine (ThermoFisher Scientific, #25030024). All cells were maintained in a humidified incubator (Panasonic) at 37°C and 5% CO_2_. Cells were passaged when 80% confluent (approximately every 2 days).

### Drug and recombinant protein treatment

Cells were treated with drugs and recombinant proteins listed in **Table 1** with the indicated incubation times.

**Table 1.**
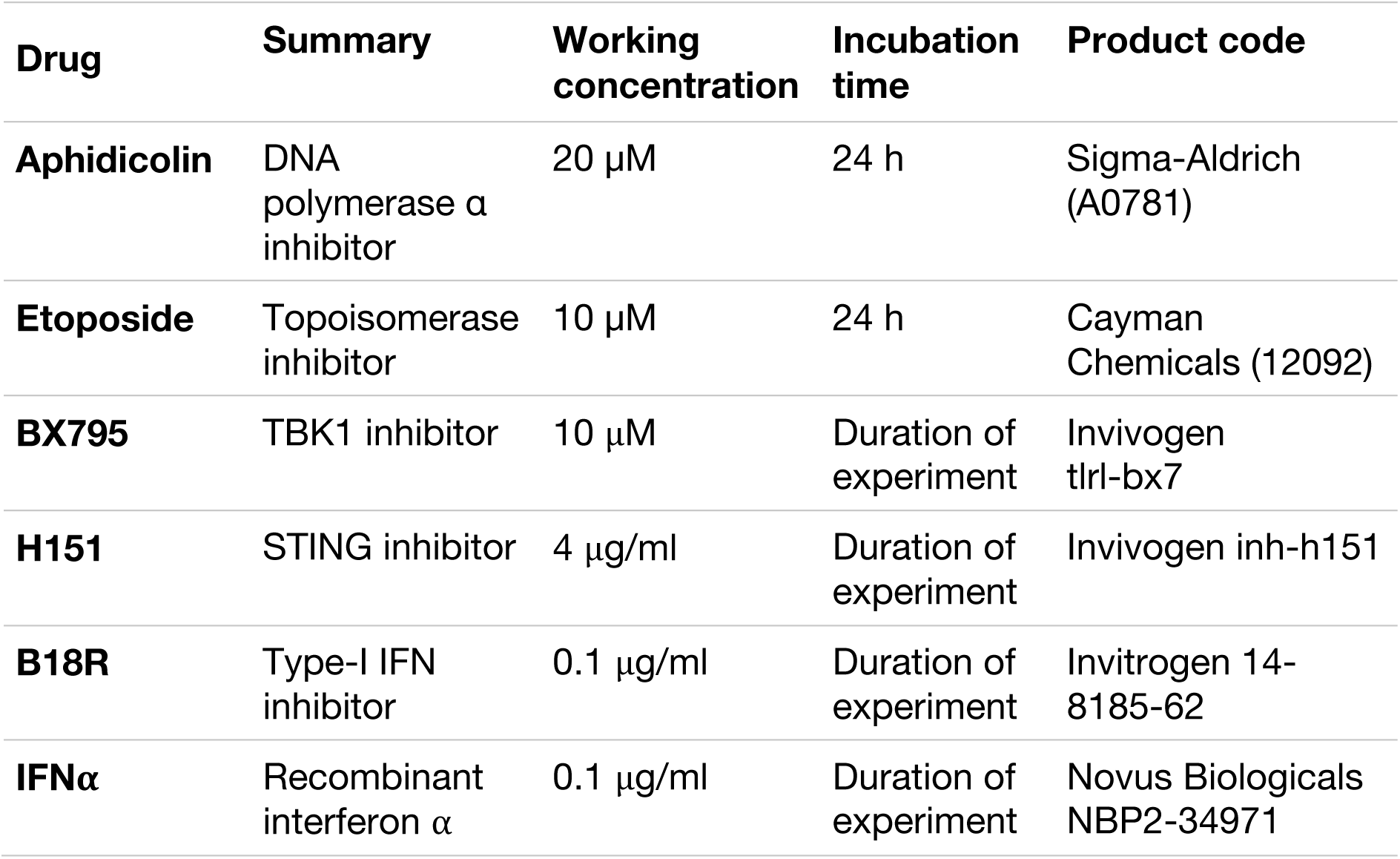
Drugs and recombinant proteins used to treat mammalian cells.

### siRNA transfection

For siRNA transfection, cells were transfected in 6 well plates by addition of 3 μl/well Lipofectamine RNAiMax (Invitrogen, 13778-150) and 20 nM of human USP18 siRNA, (Horizon SMARTpool #L-004236-00-0005), human ISG15 siRNA (Horizon SMARTpool #L-004235-03-0005), human UBE1L siRNA (Horizon SMARTpool #L-019759-00-0005) or non-targeting human siRNA (Horizon SMARTpool #D-001810-10-05) in 100 μl of serum-free, antibiotic-free DMEM. Cells were incubated for 48h before any further treatments. For transfection with mammalian expression vectors, 6 μl/well FuGENE transfection reagent (Promega) was used. pCAGGS-5HA-mISG15 was a gift from Dong-Er Zhang (Addgene plasmid # 12444) (Kim *et al*., 2004).

### Salmonella infection

Wild-type *Salmonella enterica* serovar Javiana (isolate S5-0395) and the isogenic null mutant ΔcdtB (isolate M8-0540) were kind gifts from Prof. Martin Weidmann (New York). A *Salmonella* Javiana colony was incubated in liquid LB-broth at 37°C in a shaking incubator until OD_600_ was approximately 1.0. The number of *Salmonella* at an OD^600^ of 1.0 was estimated to be 8×10^8^ bacteria/ml. Unless otherwise stated, infections were performed with an MOI of 10. The plate was centrifuged for 1 min at 1000 × g and incubated for 30 min at 37°C 5% CO_2_. Infection media was removed, cells were washed with PBS, and fresh media added containing 50 μg/ml gentamicin (Chem Cruz, sc203334) for 90 mins. This media was then replaced with fresh media containing 10 μg/ml gentamicin for the duration of the experiment.

For CFU counts, cells were washed 3× with PBS and lysed with 1% Triton X-100 in deionised sterile water for 10 mins. The supernatant was transferred to a 96-well plate, and serially diluted 10-fold. 5μl of each *Salmonella* dilution was then cultured on agar overnight in a dry incubator at 37°C. Colonies were counted manually and normalised to the 2h timepoint to calculate *Salmonella* colony forming units (CFUs).

### Microarray

Microarray samples were prepared using a standard intoxication assay of HT1080 cells with 5 ng/ml of WT- and HQ-toxin and a 48h chase. For serum-starved samples, cells were grown in media containing all components apart from 10% FBS for the full duration of one replication cycle (24 h in HT1080s). Samples were analysed at SITraN, University of Sheffield, using a human Clariom^TM^S assay (ThermoFisher Scientific, 902927). Differential expression analysis was performed with Transcriptome Analysis Console 4.0 software (Applied Biosystems, Thermo Fisher Scientific). Gene set enrichment analysis was performed using the Web-based Gene Set Analysis Toolkit (WebGestalt 2019) (Liao *et al*., 2019), using the biological process non-redundant functional database. The data is available at ArrayExpress (accession number: E-MTAB-15046).

### qRT-PCR

Standard intoxication assays were performed in HT1080 cells, and RNA was isolated using the illustra RNAspin Mini kit (GE healthcare) according to manufacturer’s instructions with 1 unit (0.5 μl) of DNase 1 (M0303S, New England BioLabs). RNA concentration was measured using a Nanodrop Lite spectrophotometer (ND-LITE-PR, Thermo Fisher Scientific) and RNA integrity tested by running the sample on a TAE agarose gel.

cDNA synthesis was performed using the RevertAid H Minus First Strand cDNA synthesis kit (Thermo scientific). qRT-PCR was performed using Maxima SYBR Green/ROX qPCR Master mix (Thermo Scientific) on the CFX96 Real-Time System. 10 μl of master mix were used per sample, and primers were added at a concentration of 0.25 μM (**Table 2**). Analysis was performed using the comparative CT Method (ΔΔ CT Method) according to guidelines by Thermo Fisher (Applied Biosystems, 2008).

**Table 2.**
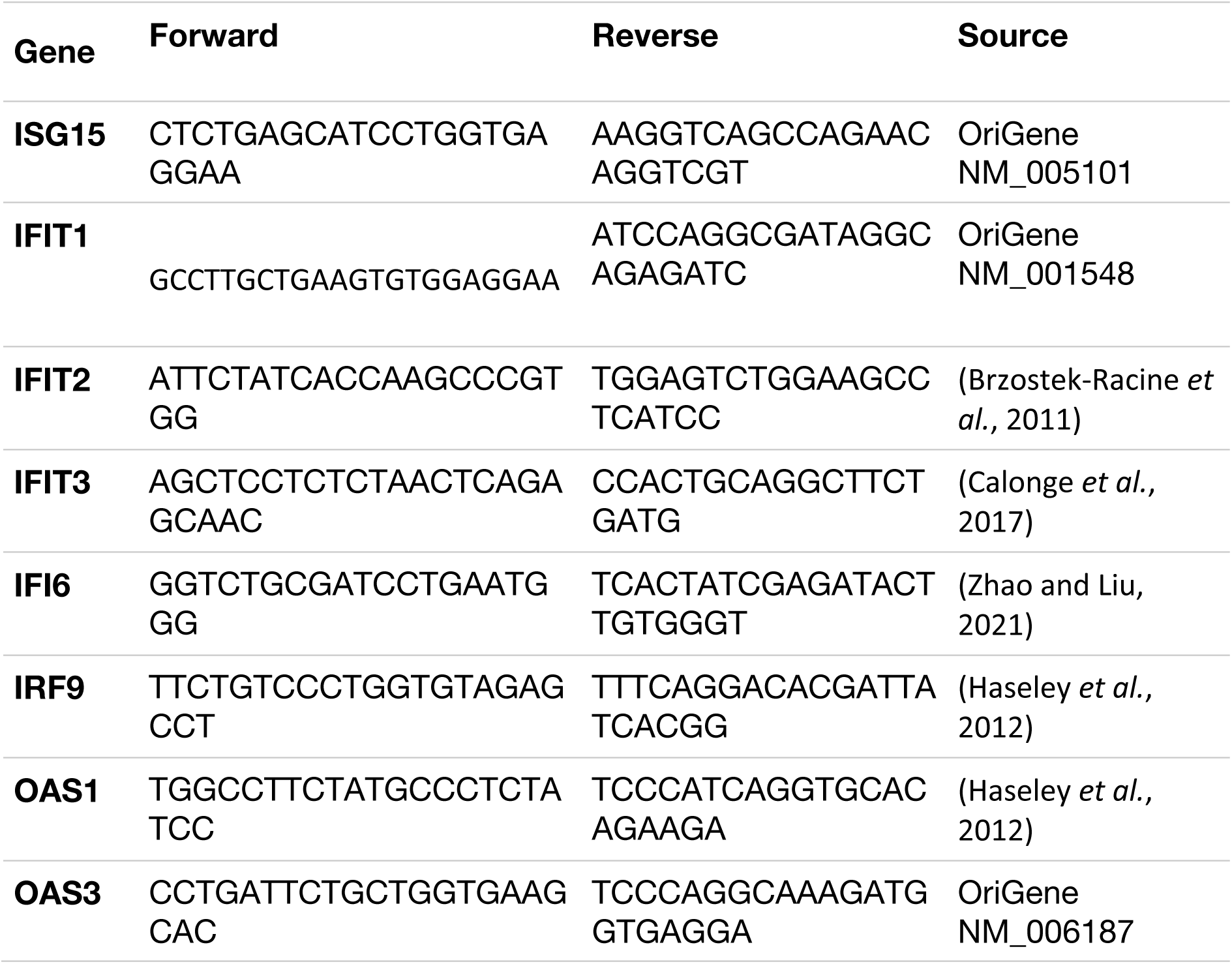
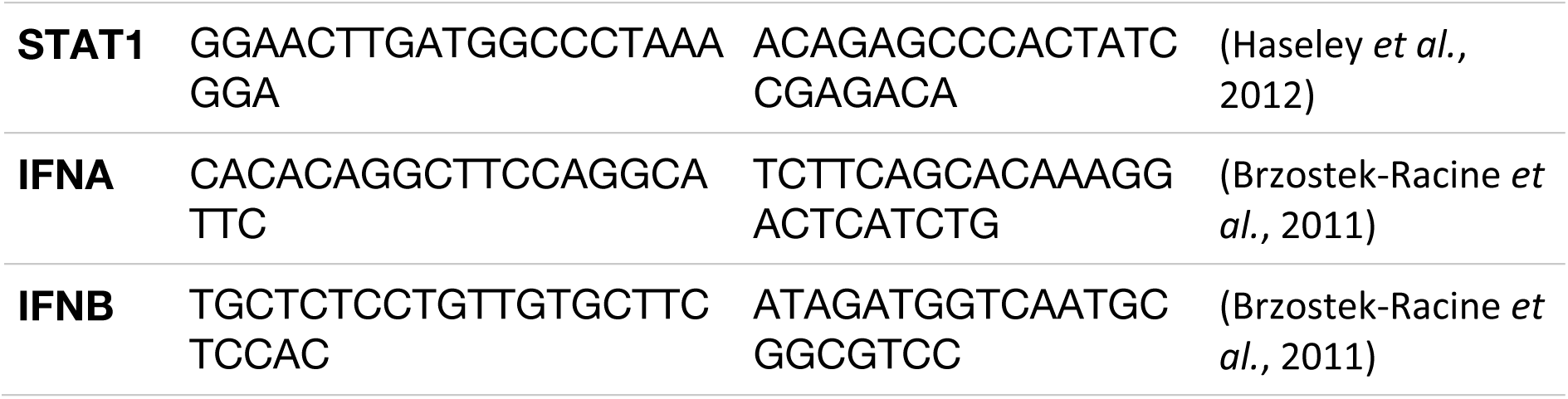
Primers used for qRT-PCR.

### Immunoblotting

Cell pellets were resuspended in sample buffer (50 mM Tris pH 6.8, 8 M Urea, 2% SDS, 0.3% Bromo blue, 1% β-mercaptoethanol). SDS-PAGE was performed using 9% Bis-tris acrylamide gels run at 40 mA/gel in Mini-PROTEAN Tetra System (Bio-Rad) in MES buffer (Life Technologies, NP0001). Transfer to PVDF membranes was performed at 400 mA for 80 min on ice or 20 mV at room temperature overnight. Membranes were blocked with 5% non-fat dried milk in TBS for 1 h at room temperature with agitation. Primary antibodies (**Table 3**) were diluted in TBS with 0.1% Tween was added overnight at 4°C with agitation. IRDye 800CW anti-mouse (926-32212, Li-Corr) and IRDye 680CW anti-rabbit (926-68073, Li-Corr) secondary antibodies were diluted in TBS with 0.1% Tween at 1:10000 for 1 hour at room temperature with agitation. Membranes were imaged at 200 μm resolution using an OdysseySa Li-Cor scanner and images processed using ImageStudioLite v5.2.5.

**Table 3.** Primary antibodies used for immunoblotting.

| **Antibody** | **Species** | **Product code** | **Dilution** |
| --- | --- | --- | --- |
| **ISG15** | mouse monoclonal | Santa Cruz sc-166755 | 1:100 |
| **Tubulin** | mouse monoclonal | Abcam ab7291 Invitrogen<br>62204 | 1:1000-1:5000 |
| **γH2AX** | rabbit monoclonal<br>mouse monoclonal | Cell Signalling 9718<br>Merck/Sigma 05-636 | 1:1000 |
| **p-pRb** | mouse monoclonal | Cell Signalling 9309 | 1:2000 |
| **Akt** | mouse monoclonal | Cell Signalling 2920 | 1:2000 |
| **p-Akt** | rabbit polyclonal | Cell Signalling 9275 | 1:1000 |
| **GAPDH** | rabbit monoclonal | Cell Signalling 2118 | 1:1000 |
| **Actin** | rabbit polyclonal | Sigma A2066 | 1:1000 |
| **p21** | rabbit monoclonal | Cell Signalling 2947 | 1:1000 |
| **USP18** | rabbit monoclonal | Cell Signalling 4813 | 1:1000 |
| **HSP90** | mouse monoclonal | Novus Biologicals NB100-1972 | 1:1000 |
| **GAPDH** | mouse monoclonal | Santa Cruz sc-365062 | 1:1000 |
| **STING** | mouse monoclonal | Proteintech 66680-1 | 1:1000 |

### Immunofluorescence

Cells were seeded in 24 well plates on glass coverslips. For staining of RPA32 pT21, cells were pre-permeabilised with PBS + 0.1% Tween for 1 minute on ice. Cells were fixed with 4% paraformaldehyde (PFA) in PBS for 10-15 mins at room temperature. PFA was removed and cells washed twice more with PBS before being stored in PBS at 4°C until staining.

For EdU staining, cells were incubated for 2h with 10 µM EdU prior to fixation. Cells were stained for EdU using a Click-iT^TM^ EdU Cell Proliferation kit with Alexa Fluor^TM^ 647 dye (ThermoFisher, C10340) as per the manufacturer’s instruction.

Cells were blocked using 3% BSA (Sigma-Aldrich, 1073508600) and 0.2% Triton X-100 (VWR, 28817.295) in PBS at room temperature for 1h. Primary and secondary antibody dilutions were prepared in PBS 0.2% Triton X-100. Primary antibodies were diluted as appropriate (**Table 4**) and added for an hour at room temperature.

**Table 4.** Primary antibodies used for immunofluorescence.

| **Antibody** | **Species** | **Product code** | **Dilution** |
| --- | --- | --- | --- |
| **ISG15** | mouse monoclonal | Santa Cruz sc-166755 | 1:100 |
| **γH2AX** | mouse monoclonal | Merck/Sigma 05-636 | 1:1000 |
| **RPA32 pT21** | rabbit polyclonal | Abcam ab61065 | 1:1000 |
| **53BP1** | mouse monoclonal | Merck MAB3802 | 1:500 |
| ***Salmonella*** | rabbit polyclonal | Abcam ab35156 | 1:1000 |

Secondary antibodies (**Table 5**) were added at a 1:500 dilution in PBS with 0.2% Triton X-100 for 30 mins at room temperature. Coverslips were then mounted on 6 μl of VectaShield mounting agent including DAPI (Vector Lab, H1200), sealed and left to dry before being imaged.

**Table 5.**
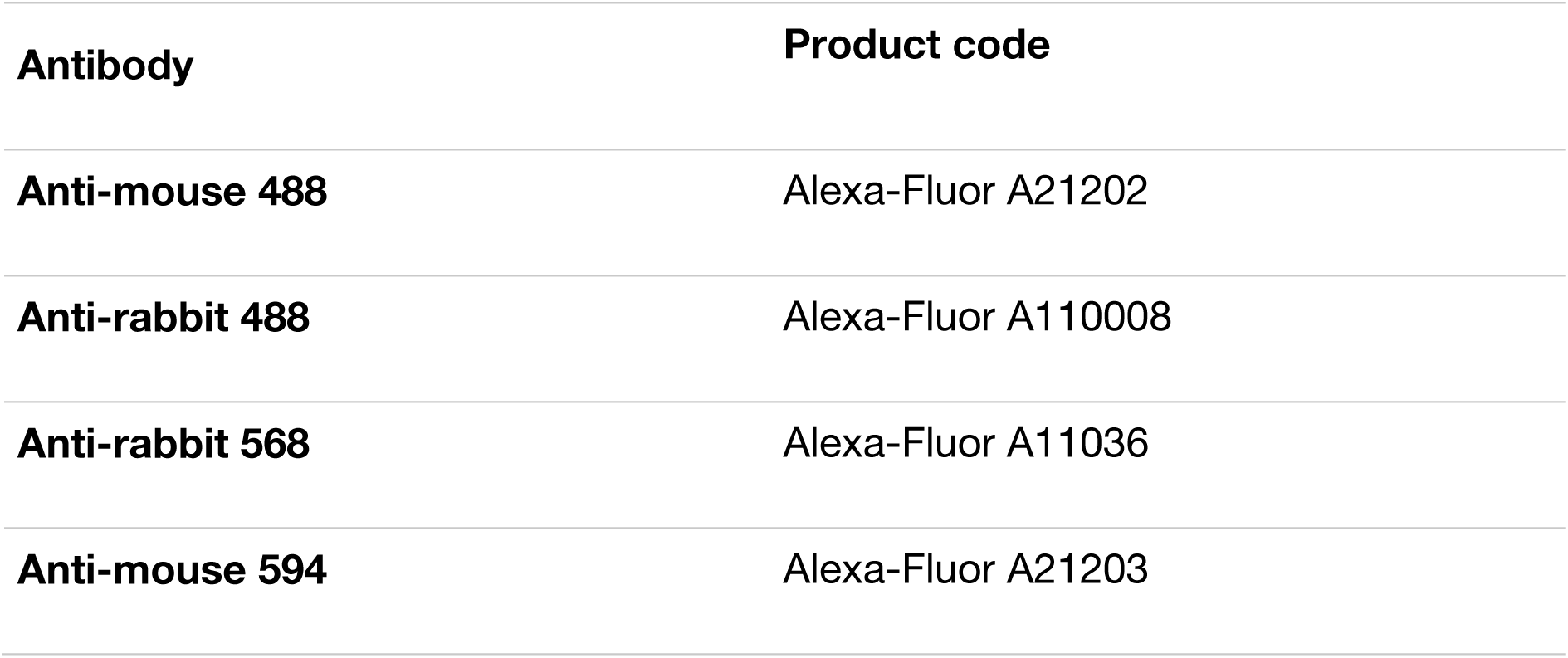
Secondary antibodies used for immunofluorescence.

| **Antibody** | **Product code** |
| --- | --- |
| **Anti-mouse 488** | Alexa-Fluor A21202 |
| **Anti-rabbit 488** | Alexa-Fluor A110008 |
| **Anti-rabbit 568** | Alexa-Fluor A11036 |
| **Anti-mouse 594** | Alexa-Fluor A21203 |

Immunostained cells were imaged using a Nikon Widefield microscope equipped with an sCMOS Andor Zyla camera. NIS elements software was used for imaging. Quantitative analysis of image intensity was carried out using the RING-tracking MATLAB code published in Ibler et al., 2019.

### Image Processing

Images were processed using Fiji v2.3. For chosen representative images, brightness and contrast were normalised, scale bar added and images cropped if necessary. The DAPI channel was converted into a binary image to obtain DAPI outlines, which were overlaid with other channels. Adobe Illustrator was used to prepare illustrations and assemble all results figures.

### Statistical analysis

Graphs and statistical analysis were carried out using GraphPad Prism 9. Generally, one-way ANOVAs were used with Šidák’s or Tukey’s multiple comparisons tests as appropriate. Significance is denoted by asterisks where *p<0.05, **p<0.01, ***p<0.001 and ****p<0.0001.

## Acknowledgements

The research was funded by a UKRI Future Leaders Fellowship to DH (MR/S034390/1, MR/X02329X/1), and CS MR/X024040/1, and supported by Medical Research Council [MR/N013840/1] PhD studentships to DS, MK. L.R. is supported by NIGMS R35GM137961. Microscopy was performed at the Wolfson Light Microscopy Facility using the Nikon Widefield system (MR/K015753/1).. Thank you to Dr. Paul Heath for assistance with the microarray experiments at the University of Sheffield.

---

**Supp. Fig. 1.**
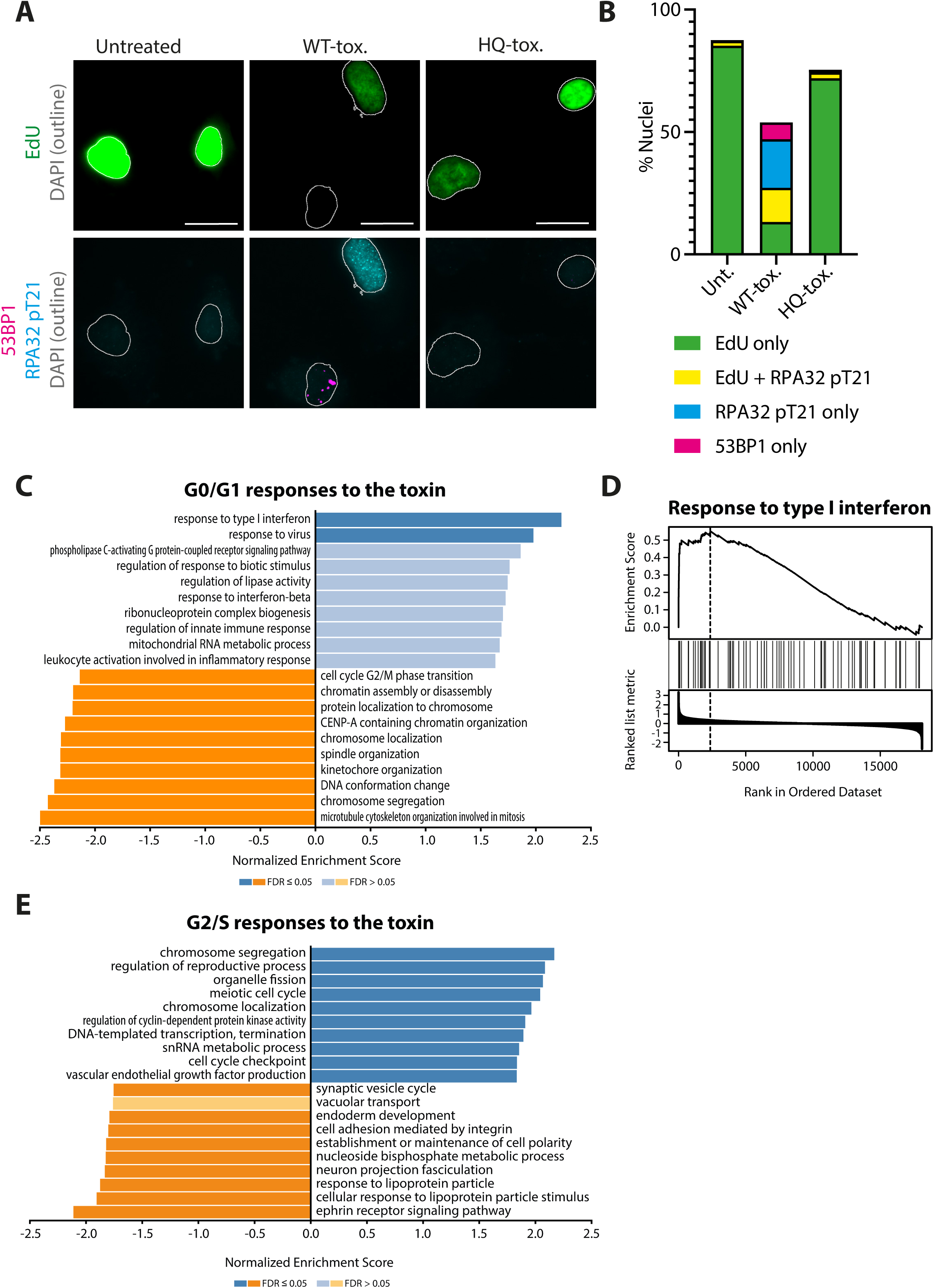
Differential gene expression in divergent responses to the typhoid toxin. **A)** Divergent DNA damage responses induced by WT- and HQ-toxin preparations were assayed in HT1080 cells using immunofluorescence of phosphorylated RPA32 (RPA32 pT21, cyan), 53BP1 (magenta) and EdU (green). Scale bars indicate 20μm. **B)** Quantification of positive nuclei in (A). Nuclei were counted as positive for RPA32 pT21 and 53BP1 if greater than the upper quartile of untreated intensity, and for EdU if greater than the lower quartile of HQ-tox intensity. Bars indicate total percentage from a single experiment with three technical replicates. **C**) Bar chart showing the top 10 positively and negatively enriched biological processes found using gene set enrichment analysis (GSEA) of 19460 genes analysed by microarray, compared between WT- and HQ-toxin treatment in serum-free media where only DNA damage responses in G0 or G1 are permissive. **D**) GSEA curve for the ‘response to type-I IFN’ biological process found in C, with the dashed line indicating the leading-edge cut-off. **E**) Bar chart showing the top 10 positively and negatively enriched biological processes found using gene set enrichment analysis (GSEA) of 19460 genes analysed by microarray, compared between WT-toxin treatment in normal and serum-free media and thus showing responses to the toxin in G2/S phase.

**Supp. Fig. 2.**
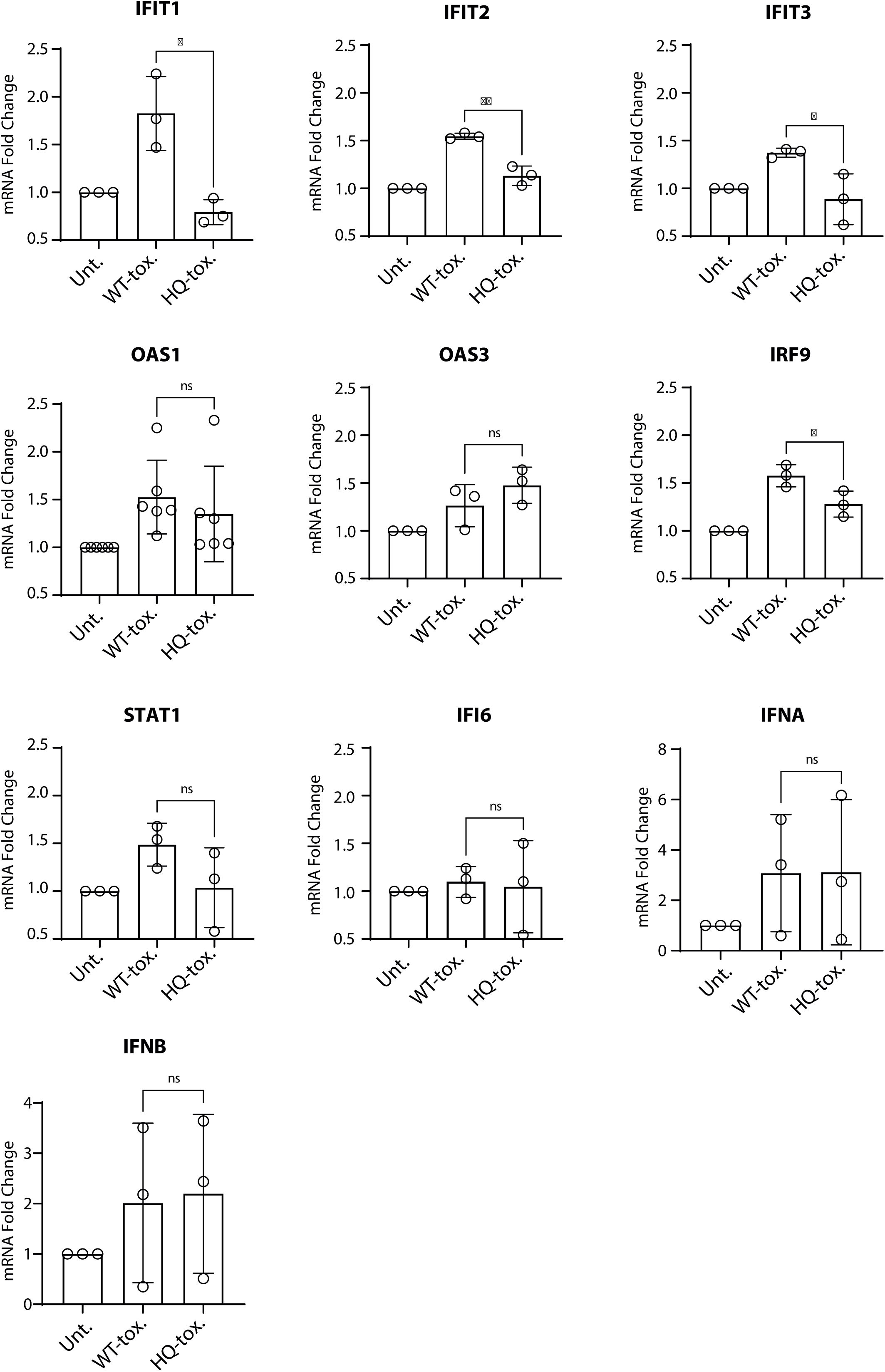
Validation of the toxin-dependent type-I interferon-like response by qRT-PCR. (**A-J**) Quantification of mRNA levels in HT1080s of 10 type-1 interferon-related genes compared between untreated, WT-toxin treated, and HQ-toxin treated samples. Bars represent mean and error bars indicate SD. Each circle is an individual RNA sample with three technical replicates. WT- and HQ-toxin conditions compared using an unpaired t-test. Asterisks indicate significance.

**Supp. Fig. 3.**
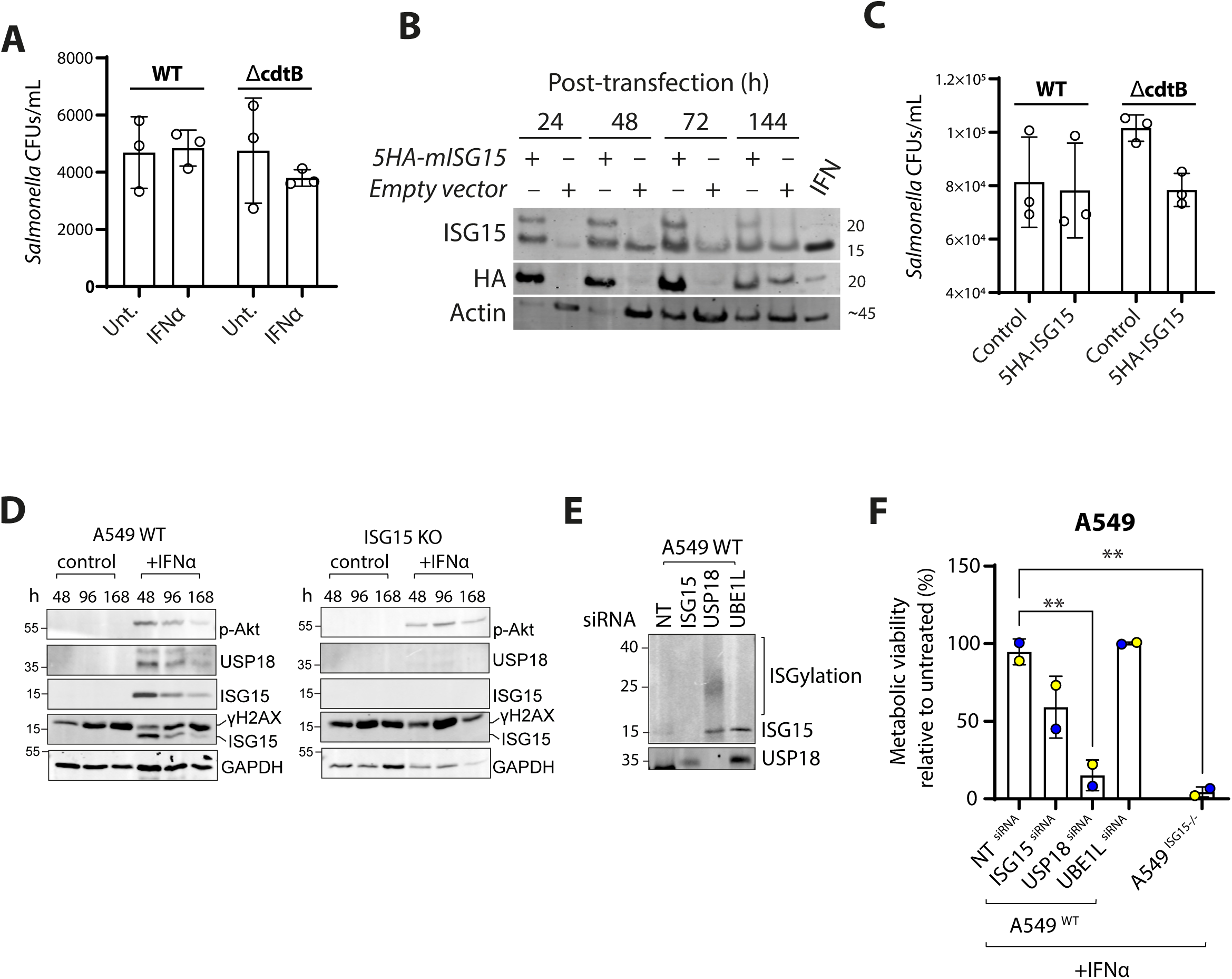
ISG15 inhibits *Salmonella* infection and is essential for cell survival in response to IFN. **A)** *S*. Javiana^WT^ and *S*. Javiana^ΔcdtB^ CFUs 24 h after infection in HT1080s with and without 72 h continuous pre-treatment with purified interferon α. Circles indicate technical repeats of a single experiment. **B**) Immunoblot of ISG15 and HA protein levels in HT1080s 24, 48, 72 and 144 h after transfection with either pCAGGS-5HA-mISG15 or empty vector pcDNA. **C**) *S*. Javiana^WT^ and *S*. Javiana^ΔcdtB^ CFUs 24 h after infection in HT1080s with and without 24 h transfection with pCAGGS-5HAmISG15 or empty vector pcDNA. Circles indicate technical repeats of a single experiment. **D)** Immunoblot of wild-type and ISG15 knockout (KO) A549 cells in the presence or absence of IFNα at 48h, 96h or 168h. MW in kDa indicated left and antibodies indicated right. **E**) Immunoblot showing ISG15 and USP18 protein levels in wild-type A549 cells 48h post-transfection with siRNA for ISG15, USP18 and UBE1L, as well as a non-transfecting (NT) control. **F**) Cell viability determined by MTT assay in wild-type A549 cells with purified IFNα treatment and transfected with siRNA for ISG15, USP18 and UBE1L. A549^ISG15-/-^ with IFN treatment is also included as a positive control for cell death. Viability is shown as a percentage of metabolic viability in the untreated control. Circles indicate independent experiments. Ordinary one-way ANOVA was used with Tukey’s multiple comparison test. Bars represent mean, error bars indicate SD, and asterisks indicate significance.

